# Budding yeast chromatin is dispersed in a crowded nucleoplasm *in vivo*

**DOI:** 10.1101/056465

**Authors:** Chen Chen, Hong Hwa Lim, Jian Shi, Sachiko Tamura, Kazuhiro Maeshima, Uttam Surana, Lu Gan

**Affiliations:** Department of Biological Sciences and Centre for BioImaging Sciences, National University of Singapore, Singapore 117543; Institute of Molecular and Cell Biology, Agency for Science Technology and Research, Proteos, 61 Biopolis Drive, Singapore 138673; Bioprocessing Technology Institute, 20 Biopolis Way, Singapore 138668; National Institute of Genetics and Sokendai (Graduate University for Advanced Studies), Mishima, Shizuoka, Japan 411-8540; Department of Pharmacology, National University of Singapore, Singapore 117543

## Abstract

Chromatin organization has an important role in the regulation of eukaryotic systems. While recent studies have refined the 3-D models of chromatin organization with high resolution at the genome sequence level, little is known about how the most fundamental units of chromatin — nucleosomes — are positioned in 3-D *in vivo*. Here we have used electron cryotomography to study chromatin organization in the budding yeast *S. cerevisiae*. Direct visualization of yeast nuclear densities shows no evidence of 30-nm chromatin fibers. Aside from pre-ribosomes and spindle microtubules, few nuclear structures are larger than a tetranucleosome. Yeast chromatin does not form compact structures in interphase or mitosis and is consistent with being in an “open” configuration that is conducive to high levels of transcription. In the absence of higher-order chromatin packing, we propose that yeast can regulate its transcription using local nucleosome-nucleosome associations.

## Introduction

Eukaryotic nuclear DNA is packaged to 1/10,000^th^ of its contour length but must remain accessible to intranuclear machinery. The nucleosome is the first level of chromatin organization: 146 bp of double-stranded DNA wraps around a histone octamer that is composed of two copies each of histones H2A, H2B, H3 and H4 (Luger et al., 1997). Chromatin organization beyond the nucleosome has been intensively studied for nearly half a century. One notable traditional-electron microscopy (EM) study of purified chromatin proposed that sequential nucleosomes are arranged into compact ~30-nm-diameter helical fibers (hereon referred to as chromatin fibers because the dimensions are variable) (Finch and Klug, 1976). Further studies proposed at least two broad classes of fiber models: the one-start solenoid (Robinson et al., 2006) and the two-start zigzag (Schalch et al., 2005; Song et al., 2014). In these fiber models, the nucleosomes pack so close that the fiber takes on the appearance of a discrete particle. The majority of these chromatin studies, however, were done *in vitro* at low ionic strength, making it unclear if the resultant models reflect chromatin organization in the crowded metabolically active interior of a cell's nucleus (Maeshima et al., 2010; Maeshima et al., 2016).

While traditional EM has revealed the overall organization of purified chromatin (Finch et al., 1975; Olins and Olins, 1974), it has provided limited insights into chromatin structure *in vivo* because macromolecular structure is highly sensitive to sample-preparation parameters: buffer conditions, chemical fixation, dehydration, and heavy-metal staining (Dubochet et al., 1988; Maeshima et al., 2010). Recently, high-throughput-sequencing-based chromatin-conformation capture (3C, 5C, Hi-C, etc.; hereon abbreviated 3C) has been used as a complementary method to study chromatin structure in fixed cells (Dekker et al., 2013; Pombo and Dillon, 2015; Smallwood and Ren, 2013). These 3C approaches reveal the most probable pair-wise chromatin contacts from a population of cells. The detected contacts are distance constraints that can be used to infer 3-D chromatin models. Single-celled 3C is also possible, but the number of detected contacts is so sparse that the resultant models are limited to larger higher-order structures like topologically associating domains (Nagano et al., 2013). Due to the dynamic nature of cells, 3C models are also susceptible to potential biases in nucleosome accessibility and fixation artifacts.

Electron cryomicroscopy (cryo-EM) permits the direct visualization of macromolecular densities in a near-native state. Furthermore, cryo-EM can provide relatively “non-invasive” windows on how macromolecular complexes interact inside of organelles and cells. For example, cryo-EM studies of vitreous sections showed that in isolated chicken-erythrocyte nuclei and partially lysed starfish and sea cucumber sperm, chromatin is condensed into 30-nm fibers (Scheffer et al., 2011; Woodcock, 1994). In contrast, studies of vitreous sections of intact HeLa and CHO cells did not reveal evidence of the 30-nm chromatin fibers (Eltsov et al., 2008; McDowall et al., 1986); instead, nucleosome densities were packed in an irregular state akin to the polymer-melt-like structure model (Maeshima et al., 2014b). Electron spectroscopic imaging of mouse cells also did not reveal any chromatin fibers (Fussner et al., 2012). Electron cryotomography (cryo-ET), which allows 3-D reconstructions of cells in a life-like state (Gan and Jensen, 2012), showed that marine picoplankton chromatin is also organized like a polymer-melt (Gan et al., 2013). Despite this growing body of evidence that the chromatin fiber is not the predominant form of chromatin packing, most studies continue to assume that chromatin packs into fibers *in vivo*. This confusion is also perpetuated because very few cryo-ET studies have been done on intact model eukaryotic cells.

The budding yeast *S. cerevisiae* (hereon referred to as yeast) is an important model system for chromatin studies. Fluorescence-microscopy imaging of certain genomic loci (Bystricky et al., 2004) and high-resolution nucleosome-positioning studies (Brogaard et al., 2012) produced models of yeast chromatin that were consistent with chromatin fibers. This conclusion is controversial because 3C studies did not detect any evidence of chromatin fibers (Dekker, 2008; Hsieh et al., 2015). While light-microscopy and high-throughput sequencing-based approaches have produced important advances to our understanding of chromatin structure, no study has directly visualized nucleosomes within the crowded molecular environment of intact yeast.

In this work, we directly visualized the nuclear densities of yeast in 3-D using cryo-ET of vitreous sections. We controlled for sample-preparation artifacts using known chromatin structures. Our analysis of cryotomograms of G1-and metaphase-arrested yeast did not uncover any evidence of regular 30-nm chromatin fibers. Instead, nucleosomes have an irregular organization and do not adopt any higher-order structures. Nucleosomes do frequently pack close enough to form small clusters. Given the low frequency of introns in yeast and the nucleosome-occupancy data showing nucleosome depletion near the transcription start site (Lee et al., 2007), we propose that some of the small clusters of nucleosomes may in fact contain the coding regions of genes.

## Results

### Chromatin fibers are compact and stable

The diversity of model systems and cryo-EM techniques makes it challenging to understand what are the most reproducible structural features of chromatin. Purified chromatin fibers are thin enough that they can be plunge-frozen and immediately imaged by cryo-EM. Chromatin inside cells cannot be effectively imaged by cryo-EM unless the cells are first thinned in cryogenic conditions; cryomicrotomy can produce such vitreous sections (Dubochet et al., 1988). To account for the effects of cryo-EM sample preparation, we used purified chicken-erythrocyte chromatin fibers as a positive control (Fig. S1). We stabilized the chromatin fibers in isolation buffer containing 2 mM Mg^2+^ (Widom, 1989), both with and without cryoprotectant, and then performed cryo-ET on these samples prepared either by plunge-freezing or by cryomicrotomy (Figs. 1 and S2). Pairwise comparisons of the resultant cryotomograms lead to the following conclusions: (1) chromatin fibers are recognizable as compact particles regardless of sample-preparation technique; (2) these fibers are so compact that it is difficult to distinguish individual nucleosome densities when the fibers aggregate; (3) in cryosections, chromatin fibers remain intact and are compressed along the cutting direction, as expected. Having controlled for the technical aspects of cryo-ET samples, we next combined cryomicrotomy with automated cryo-ET to image many yeast cells to ensure that our observations were reproducible (Tables 1 and S1).

**Figure 1.**
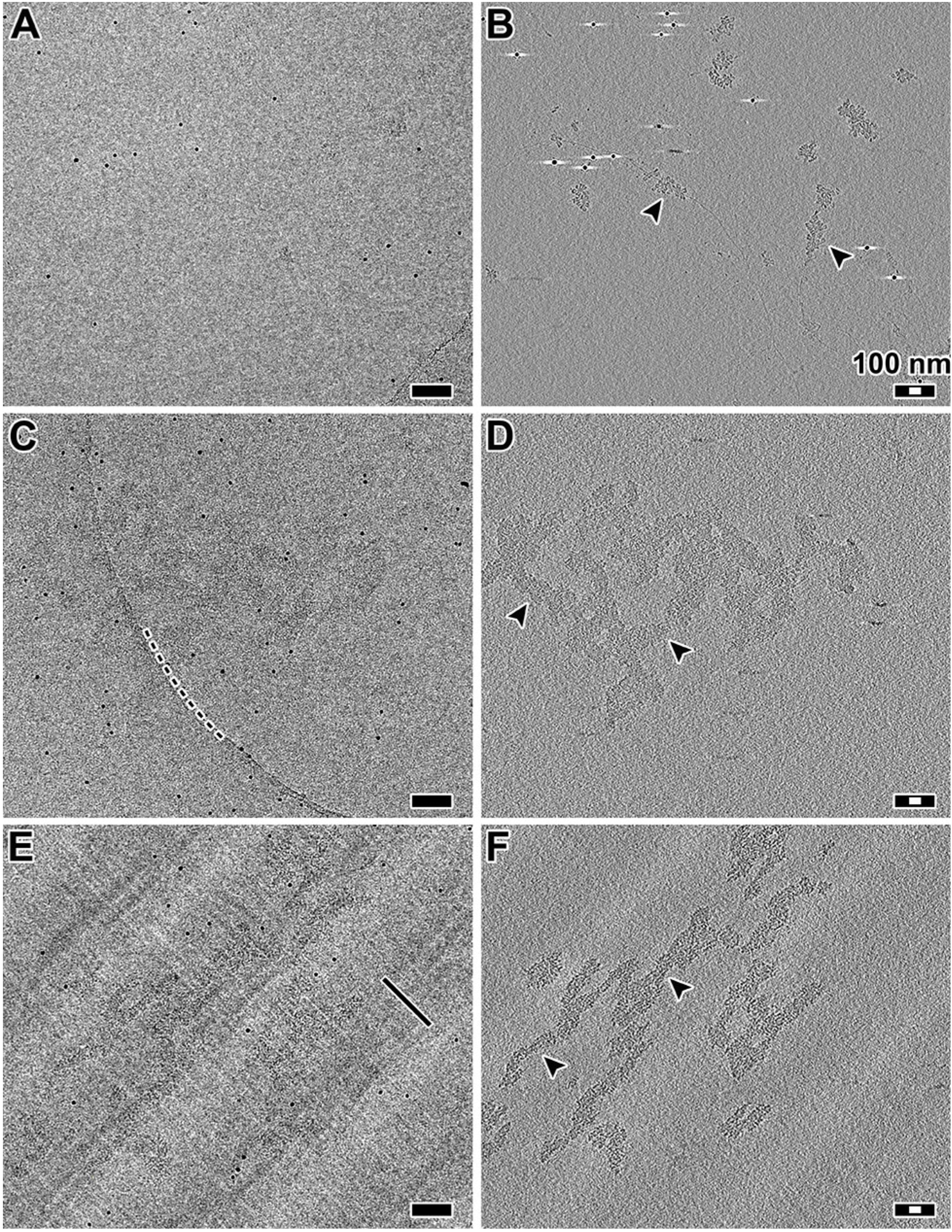
Chromatin fibers are compact and remain intact in cryosections. **(A)** Projection image of chicken erythrocyte chromatin fibers plunge-frozen in dialysis buffer. **(B)** Tomographic slice (12-nm thick) of the position in (A). Arrowheads indicate examples of chromatin fibers. **(C)** Projection image of chicken erythrocyte chromatin fibers plunge-frozen in the presence of dextran. Note that due to low dose (~2 e/Å^2^ per projection) and relatively small defocus (−6 μm), the fibers are difficult to see. The dark, punctate densities are 10-nm gold fiducials. The dashed line marks the curved edge of the holey carbon support. **(D)** Tomographic slice (12-nm thick) of the position in (C). Arrowheads indicate examples of chromatin fibers. **(E)** Projection image of a frozen-hydrated section containing chromatin fibers. Knife marks are thin linear features that are parallel to the cutting direction, as indicated by the black arrow. **(F)** Tomographic slice (12-nm thick) of the same area as in (E). Arrowheads indicate examples of chromatin fibers. The alternating light-dark background bands running from the lower left to upper right of C and D are crevasse artifacts, which are visible due to the proximity of the tomographic slice to the cryosection surface. Black scale bars are 100 nm; white scale bars are 30 nm.

**Table 1.** Summary of chromatin conformations observed.

| Sample | Treatment | Observations | Data | Notes |
| --- | --- | --- | --- | --- |
| CEN + 2 mM<br>Mg<sup>2+</sup> | PF, no Dextran | Disperse 30-nm<br>fiber | Figs. 1, S2 | + control |
| CEN + 2 mM<br>Mg<sup>2+</sup> | PF, Dextran | Aggregates of<br>fiber | Figs. 1, S2 | + control |
| CEN + 2 mM<br>Mg<sup>2+</sup> | CEMOVIS, Dextran | Aggregates,<br>compressed | Figs. 1, S2 | + control |
| Wild-type cells | CEMOVIS, fixed,<br>Dextran* | Irregular<br>chromatin | Figs. 4, S4C | + control,<br>*in vivo* |
| G1 cells | CEMOVIS, Dextran* | Irregular<br>chromatin | Figs. 2, 3, 4, 5,<br>S4A, S5 | *in vivo* |
| Metaphase cells | CEMOVIS, Dextran* | Irregular<br>chromatin | Figs. 2, 3,<br>S4B, Movie S1 | *in vivo* |

### Chromatin does not have long-range order in yeast

In many eukaryotes, chromosomes undergo global reorganization, from an “open” interphase state to a condensed mitotic state. While it remains controversial how much yeast mitotic chromatin condenses (see discussion), yeast chromatin might form more fibers in mitosis than in interphase. To test for this condensation, we arrested cells at both G1 and metaphase (Fig. S3) and then imaged them by cryo-ET of cryosections. Nucleosome-like densities were abundant inside the nuclei of both kinds of cells, but higher-order chromatin structures that resemble fibers or highly compact arrays were absent (Figs. 2A and 2B and Movie S1; 2C and 2E). In fact, we did not see any assemblies of nucleosome-sized particles that have long-range order of any kind. Ribosome-like particles - most likely pre-ribosomes (Tschochner and Hurt, 2003) — were also present in the nucleus (Figs. 2A and 2B; 2G and 2H). In addition, in metaphase cells, spindle microtubules, which have a 25-nm diameter, could be seen inside the nucleus. Visualization of intranuclear macromolecular complexes that of comparable size to chromatin fibers further demonstrates that our data has enough contrast to reveal chromatin fibers. In summary, these data show that yeast chromatin does not have features consistent with chromatin fibers or compact chromatin structures of any kind, in G1 or metaphase.

**Figure 2.**
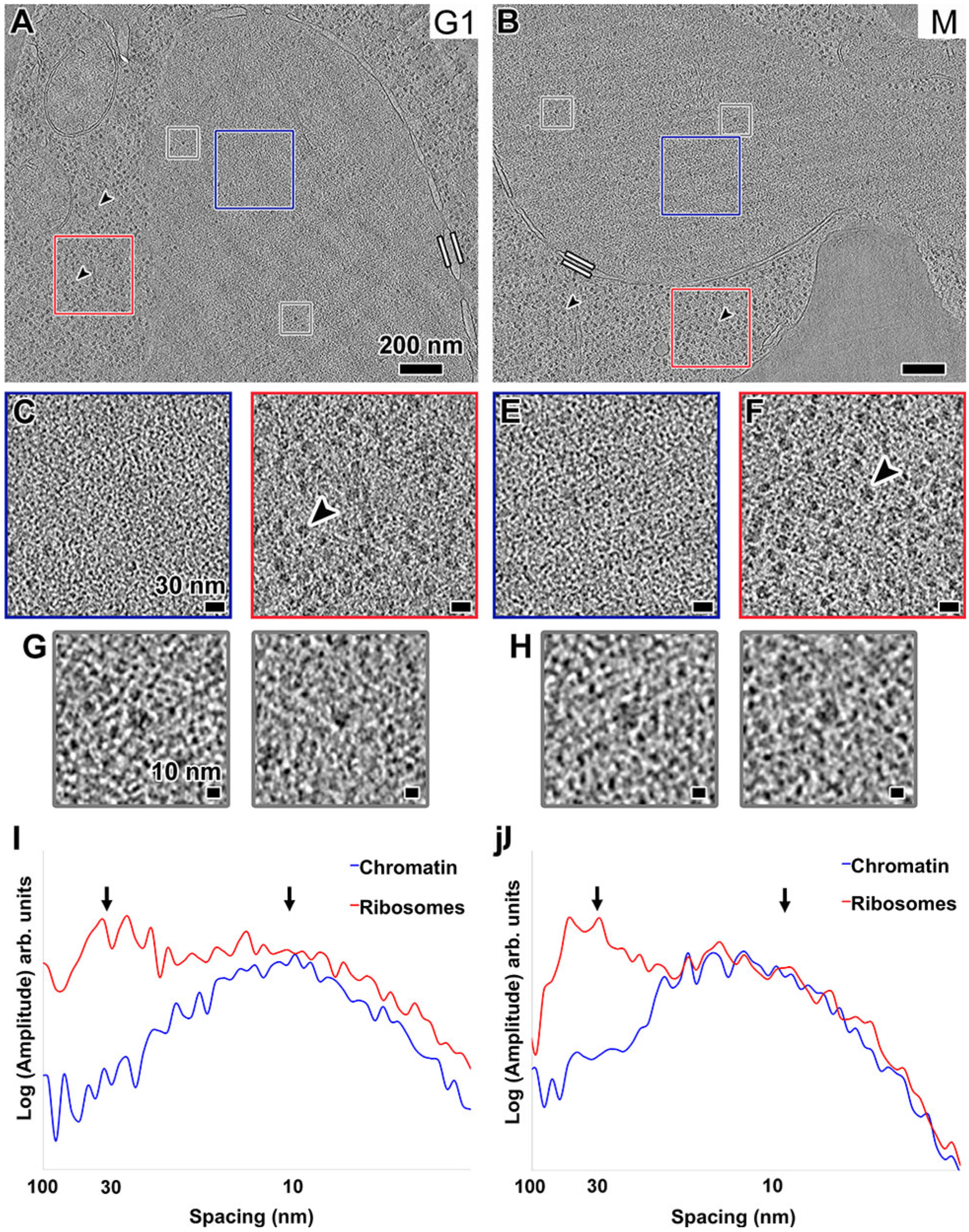
Chromatin is not organized as fibers in yeast. Tomographic slices (30-nm thick) of yeast nuclei in **(A)** G1 and **(B)** metaphase (M) cells. The nuclei (Nuc) and a mitochondria (Mi) are labeled. Parallel white bars mark inner-and outer-nuclear membranes. Black arrowheads point to cytoplasmic ribosomes. Scale bars are 200 nm. **(C)** and **(E)** Enlargements (3-fold) of the intranuclear positions enclosed by blue boxes in panels (A) and (B), respectively. **(D)** and **(F)** Enlargements (3-fold) of cytoplasmic ribosomes enclosed by red boxes in panels (A) and (B), respectively. Scale bars are 30 nm. **(G)** and **(H)** Examples of intranuclear ribosomesized densities boxed out (grey) from (A) and (B), respectively, and enlarged 6-fold. Scale bars are 10 nm. **(I)** and **(J)** Rotationally averaged power spectra of chromatin-and ribosome-rich positions from (C) and (D), (E) and (F), respectively. Arrows point to 30- and 10-nm spacings.

Fourier analysis is a well-established method to detect the presence densely packed regular particles like nucleosomes and chromatin fibers (Eltsov et al., 2008; Scheffer et al., 2011). To detect and characterize any regular motifs that may be present, we performed Fourier analysis on positions within the nuclei (Figs. 2C and 2E). A broad peak at ~10-nm spacing stood out in both G1 and metaphase cells (blue plots in Figs. 2I and 2J). This signal is expected from loosely packed nucleosomes, which are 6-nm thick and 11-nm in diameter (Joti et al., 2012; Nishino et al., 2012). In contrast, we did not observe a peak at ~30-nm spacing, which would be expected of a nucleus that is enriched with 30-nm chromatin fibers (Scheffer et al., 2011). These observations were reproducible in all 19 our cryosectioned yeast samples (see Figs. S4A and S4B for more examples; Table S1).

As an internal control, we analyzed the cytoplasms of both G1 and metaphase cells. Many of these positions are densely packed with ribosomes (Figs. 2D and 2F), which produced the expected broad peak at ~ 30-nm spacing (red plots in Figs. 2I and 2J). To eliminate even the remotest possibility that the effects of microscope underfocus conditions caused us to miss the chromatin fibers, we also acquired several tilt series much closer to focus (Fig. S5). This close-to-focus data did not show any evidence of chromatin fibers.

The high contrast in our best tomograms allowed us to render the nuclear volumes as isosurfaces so that the nucleosome-like densities could be visualized in 3-D (Fig. 3). This rendering style enables the direct inspection for 3-D arrangements of chromatin structures that we might otherwise have missed when inspecting 2-D “tomographic” slices. As expected, the isosurfaces confirmed the crowded and irregular nature of the yeast nuclear structures (Figs. 3B, C and 3F, G). This crowdedness was even more evident when we increased the thickness to 70 nm (Figs. 3D and 3H). In summary, our cryo-ET data show no evidence that yeast chromatin organizes as fibers or any periodic higher-order structures *in vivo*.

**Figure 3.**
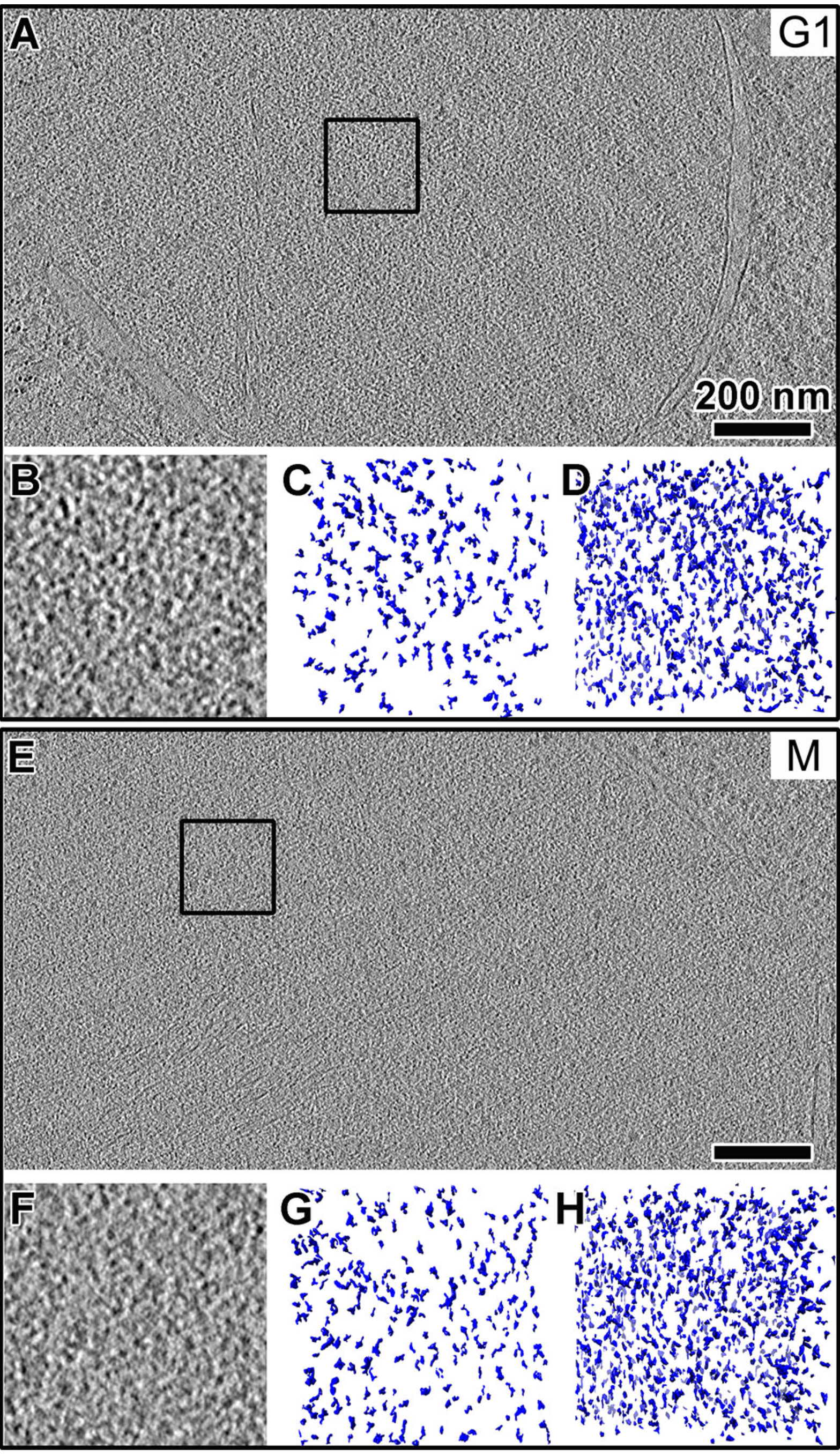
Yeast nuclei are crowded, but do not have highly ordered chromatin complexes. Tomographic slices (10-nm thick) of nuclei in a **(A)** G1 and **(E)** metaphase (M) cell. Scale bars are 200 nm. **(B)** and **(F)** Enlargements (3-fold) of the intranuclear positions enclosed by boxes in panels (A) and (E), respectively. **(C)** and **(G)** Isosurface rendering of a 10-nm-thick volume of the region in (B) and (F), respectively. Note that some of the smaller densities are from tomographic slices just “above” and “below” the selected volume. **(D)** and **(H)** Isosurface rendering of a 70-nm-thick volume centered on the same region in (B) and (F), respectively.

### Fixed cells also have disorganized chromatin

A recent study used the new 3C variant called “Micro-C” to study chromatin structure in formaldehyde-fixed yeast cells (Hsieh et al., 2015). That study concluded that yeast chromatin does not form chromatin fibers but instead packs into tetranucleosome clusters. Since 3C approaches are thought to capture native chromatin interactions, they could inform on our observations if the formaldehyde-fixation step does not seriously disrupt nuclear structure. We therefore tested if fixed cells are seriously perturbed and if they have the proposed oligonucleosome structures. We fixed log-phase wild-type cells (US1363) in formaldehyde using published protocols (Hsieh et al., 2015) and then high-pressure froze, cryosectioned, imaged, and generated cryotomograms using the same conditions as for unfixed cells. Overall, we did not see any gross distortions to cellular morphology except to the mitochondrial membranes (Fig. 4). Macromolecular complexes such as ribosomes did not form aggregates either. While the contrast was not as high as in unfixed cells, we could see that the nucleosome-like densities were still organized irregularly (Fig. 4C). Some nucleosome-like densities were also close enough to form contacts (Fig. 4D), but we did not see any highly compacted structures reminiscent of tetranucleosomes in crystals (Schalch et al., 2005) or in glutaraldehyde-fixed chromatin arrays (Song et al., 2014). These observations were reproducible in all our fixed yeast samples (see Fig. S4C).

**Figure 4.**
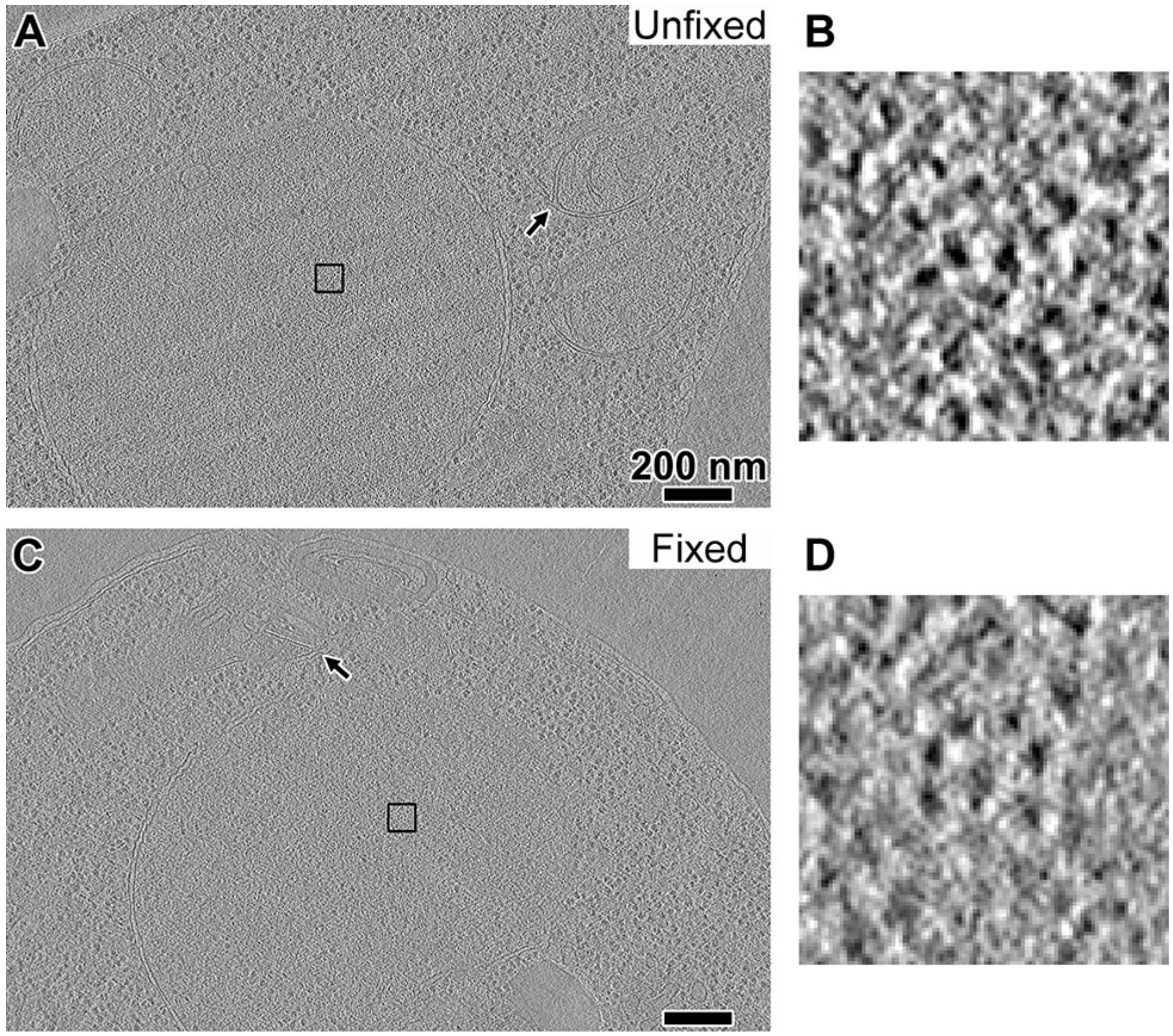
Fixation does not seriously perturb nuclear structure. **(A)** A tomographic slice (10-nm thick) of an unfixed US1363 G1-arrested cell; **(B)** A 15-fold enlargement of the nuclear densities boxed in (A). **(C)** A tomographic slice (10-nm thick) of a formaldehyde-fixed US1363 cell. **(D)** A 15-fold enlargement of the nuclear densities boxed in in (C). Arrows point to the mitochondrial membranes. Scale bars are 200 nm.

### Local chromatin structure in yeast

If groups of nucleosomes form abundant, well-defined complexes, they must also appear frequently as clusters of intranuclear densities in our cryotomograms. This notion has been used in the study of chromatin fibers in chicken erythrocytes (Scheffer et al., 2011), ATPases in mitochondria (Davies et al., 2011), and polysomes in *E. coli* and neurons (Brandt et al., 2010; Brandt et al., 2009). We could clearly see small clusters of nucleosome-like densities in many of our best cryotomograms. These clusters may, for example, be tetranucleosomes, albeit not as densely packed as in the aforementioned crystal and cryo-EM structures (Robinson et al., 2006; Schalch et al., 2005; Song et al., 2014). To locate these candidate oligonucleosome structures in a more objective and automated way, we searched using template matching (Fig. 5A). Not knowing how exactly the nucleosomes would be positioned next to each other, we used as references simple clusters of spheres packed to ~10-nm center-to-center distance (Fig. 5B). Very few template-matching hits looked exactly like the reference models, further supporting our observation that yeast chromatin is irregular. Of the best-correlating hits, the positions of the nucleosome-like densities varied so much that further analysis by subtomogram averaging was not feasible (Fig. 5B). Furthermore, fewer than ~10 clusters of each “class” could be found in the search volume. If we extrapolated to the entire yeast nucleus, then there would be fewer than ~1,000 of each of these classes of oligonucleosome clusters. While it was tempting to template match with other arrangements of closely packed spheres, many of these would yield overlapping hits because the clusters share similar motifs (like two nucleosomes in a row). This analysis shows that while clusters of nucleosomes exist in yeast, they are arranged in many different configurations.

**Figure 5.**
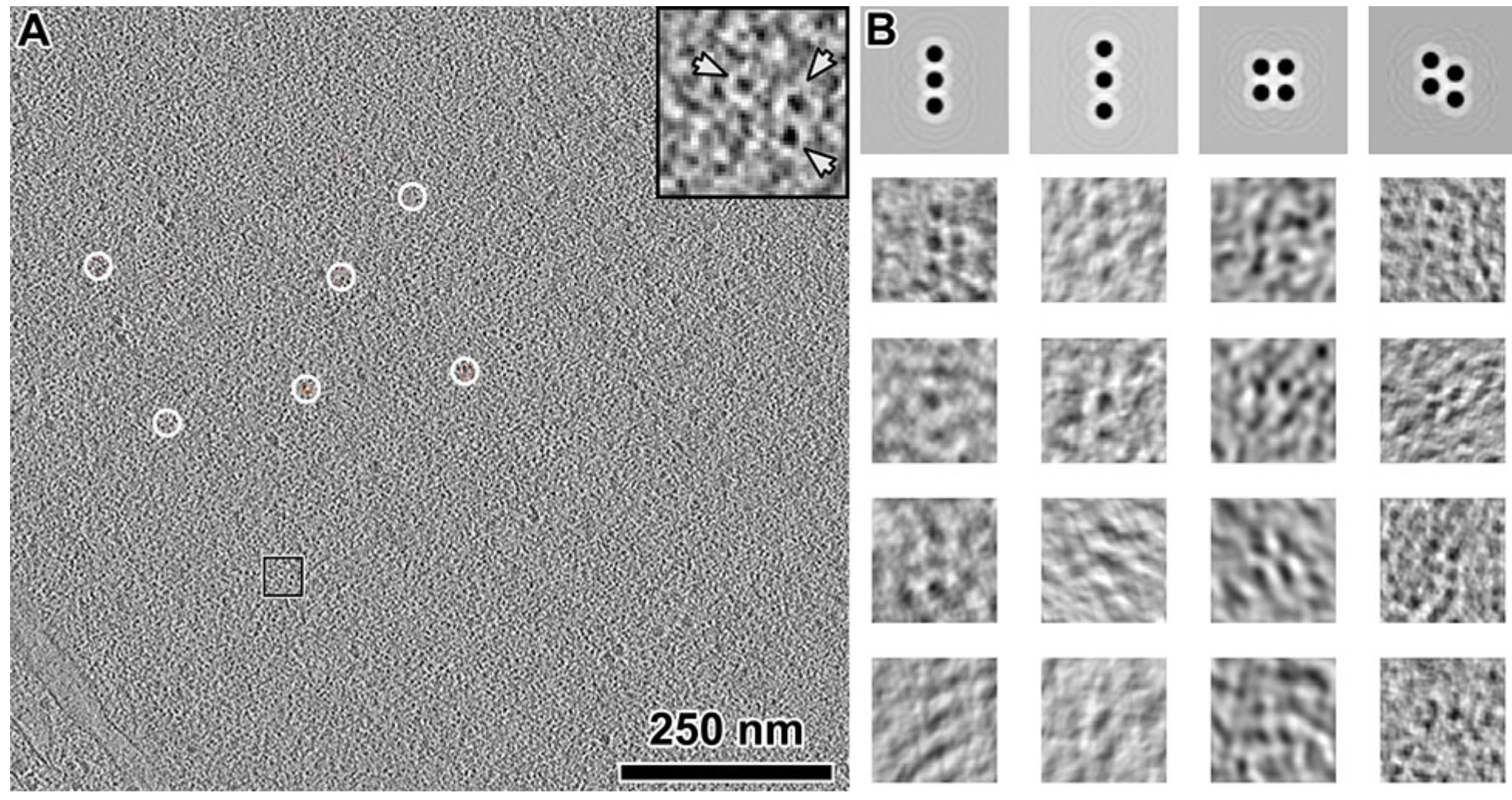
Oligonucleosome-like densities are heterogeneous. **(A)** A tomographic slice (10-nm thick) of the interior of a G1 nucleus. The few template matching hits of a candidate tetranucleosomes are circled, but only part of the density is within this 10-nm-thick volume. **(B)** Upper row: Four different templates showing potential clustering of nucleosomes, enlarged 6-fold relative to (A). Lower rows: examples of extracted and aligned template-matching hits (10-nm thick tomographic slice), showing the heterogeneous nature of these particles. The missing wedge causes some of the densities to appear elongated along one direction.

## Discussion

### Vitreous sections reveal details of *in vivo* chromatin organization

Cryo-ET makes it possible to address structural cell-biology problems at molecular resolution (Dubrovsky et al., 2015). Limitations imposed by electron-scattering physics nevertheless have restricted the vast majority of such advances to bacteria, which are thin enough to be plunge-frozen and then imaged *in toto* (Pilhofer and Jensen, 2013). Cryo-ET studies of eukaryotes, particularly the investigations of minimally perturbed chromatin require that intact cells be thinned in the frozen-hydrated state. This challenge can be surmounted by cryomicrotomy, which can preserve molecular details in a near-native state (Al-Amoudi et al., 2004). We have now used cryo-ET of vitreous sections to resolve details of yeast chromatin and have revealed nucleosome-organizational principles that may inform on how chromatin organization may influence transcription.

Our study reveals that the yeast nucleus is crowded with macromolecular complexes, without any evidence of chromatin fibers or anything that could be a densely packed chromatin structure. The absence of periodic chromatin structures in yeast is reminiscent of the picoplankton *Ostreococcus tauri* (Gan et al., 2013), and HeLa cells (Mahamid et al., 2016), the only other eukaryotes studied intact by cryo-ET. More cryo-ET studies of eukaryotes are needed to establish whether chromatin fibers are the exception or the rule.

### Yeast chromatin does not form condensed structures

Yeast has been studied extensively using EM of fixed and stained cells, but there was no evidence of mitotic chromosome condensation, perhaps as a result of sample-preparation artifacts (O'Toole et al., 1999; Robinow and Marak, 1966; Winey et al., 1995). In contrast, both fluorescence *in situ* hybridization and Lac-operator-array tagging experiments have shown that distant sequences on the same chromosome move closer together during mitosis (Guacci et al., 1994; Vas et al., 2007). Cryo-EM of cryosections can reveal condensed chromosomes like those in metaphase HeLa cells because the local concentration of nucleosomes increases at positions corresponding to chromatids (Eltsov et al., 2008). In our cryotomograms of metaphase-arrested cells, we could not detect any condensed chromosomes. Therefore yeast undergoes mitotic condensation without increasing local nucleosome concentration. The simplest explanation is that yeast chromosomes condense by means of looping interactions, perhaps as proposed by a recent study (Cheng et al., 2015).

The Micro-C approach was recently developed to probe the 3C “blind spot”, making it possible to detect chromatin structures corresponding to a few nucleosomes (Hsieh et al., 2015). This study also did not find evidence for chromatin fibers in yeast. However, the mapping of Micro-C data onto a 3-D chromatin model depends on some assumptions that have not yet been controlled for (Pombo and Dillon, 2015). Perhaps the most critical factor is how much the nucleus is perturbed by the fixation step. We have now shown that formaldehyde fixation used in 3C does not seriously perturb the nucleus. Therefore, cryo-ET and 3C yield the same conclusion that there is no evidence of chromatin fibers or large-scale nucleosome assemblies in yeast *in vivo*. As cryo-ET and 3C further improve, we will gain more complementary insights into chromatin structure. Chromatin structural models of yeast can be further improved via the integration of super-resolution fluorescence *in situ* hybridization (Boettiger et al., 2016) and fluorescence microscopies (Ricci et al., 2015).

### The meaning of higher-order chromatin

The prevailing model of higher-order chromatin is hierarchical. Nucleosomes are packed as highly ordered ~30-nm helical fibers (Fig. 6A). These fibers can then fold into ~130-nm-thick coiled structures called ‘chromonema fibers’, which fold into yet-larger structures such as mitotic chromosomes (Belmont and Bruce, 1994). None of these structures have been observed in the two small eukaryotes studied intact by cryo-ET: *S. cerevisiae* and the picoplankton *O. tauri*. While most of the nucleosome densities can be explained by polymer-melt-like chromatin (Fig. 6B), some densities pack into smaller oligomers containing fewer than ~5 nucleosomes. Some of these densities appear in a linear series, perhaps analogous to the face-to-face stacking motif seen in purified starfish sperm chromatin (Scheffer et al., 2012). We therefore propose that at least in smaller eukaryotes, chromatin is regulated by “lower-order” structures.

**Figure 6.**
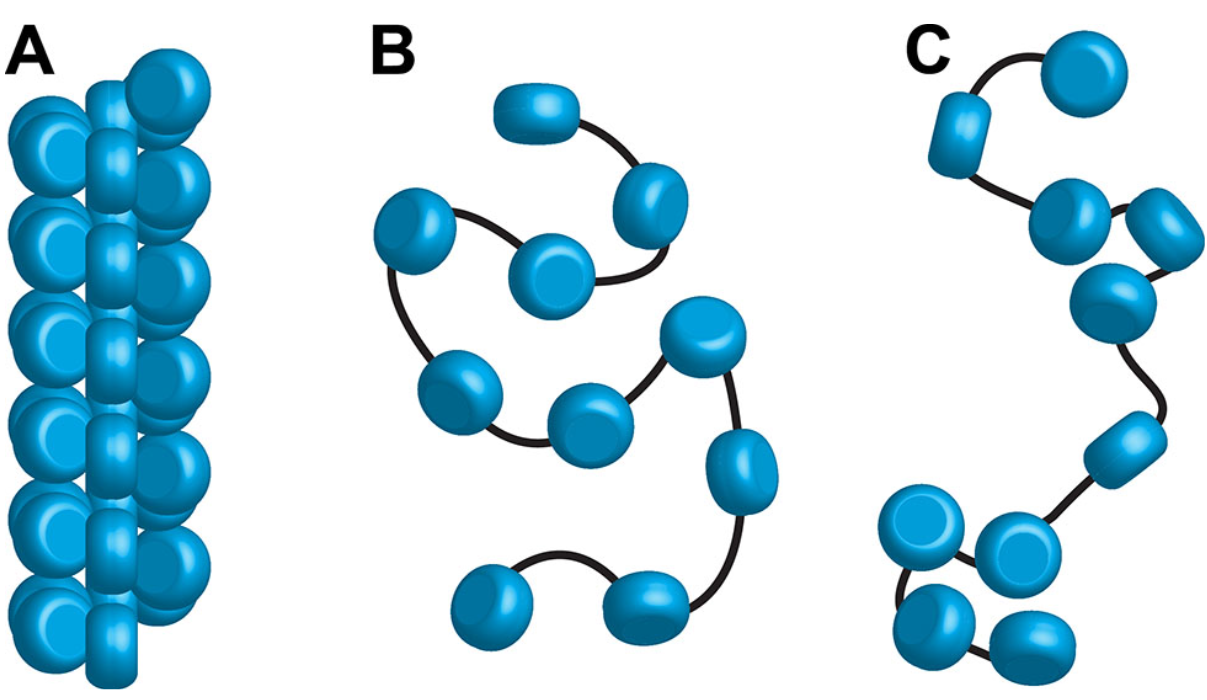
Yeast chromatin has a polymer-melt-like structure with local nucleosome clustering. Three levels of chromatin organization: **(A)** regular 30-nm fiber, **(B)** disordered polymer melt-like chromatin, and **(C)** polymer-melt interspersed with local nucleosome clusters. Our cryo-ET data is more consistent with nucleosomes (blue cylinders) packing without long-range order (B and C). Black lines indicate how linker DNA might connect adjacent nucleosomes.

*S. cerevisiae* has an ultra-high gene density and an ultra-low abundance of introns (Derelle et al., 2006), meaning that on average, the coding region of each gene spans fewer than ~10 nucleosomes. Micro-C reveals that the most probable interchromatin interactions are along the diagonal of the contact map (Hsieh et al., 2015), i.e., between sequential nucleosomes. Nucleosome-occupancy data have shown that eukaryotic genes have a nucleosome-depleted region at the transcription start site (Lee et al., 2007). The combination of these high-throughput-sequencing-based models and our direct-imaging data suggests that the coding regions of genes can fold into oligonucleosome clusters. In the absence of chromatin fibers, the overall nucleosomal arrangement is likely to be zigzag, punctuated by short stretches of extended linker DNA (Fig. 6C).

### Biological factors that disfavor higher-order chromatin

Together with the previous reports, our study raises questions about the notion of higher-order structure, at least in proliferating cells. Indeed, the only two cell types shown to have chromatin fibers — by cryo-EM — are starfish sperm and chicken erythrocytes (Scheffer et al., 2011; Woodcock, 1994), both of which are terminally differentiated cells that have minimal transcriptional activity. What factors, then, inhibit fiber formation? Yeast chromatin is highly acetylated (Clarke et al., 1993), which would destabilize the critical ionic interaction between the histone H4 N-terminal tail and the acidic patch on adjacent nucleosomes (Shogren-Knaak et al., 2006). Hence, the extent of acetylation may be important for the modulating of chromosome compaction. Another factor that may influence chromatin fiber formation is that yeast has an unconventional linker histone compared to those found in chickens (Harshman et al., 2013). It will therefore be valuable to image the chromatin of cells that have low levels of acetylation and proliferating cells that have conventional linker histones.

### Biological consequence of nuclear architecture

The absence of fibers and chromatin condensation in yeast leads to profound consequences because highly compact chromatin (Fig. 6A) exposes less sequence to transcriptional machinery than loosely packed chromatin (Figs. 6B, C). Transcriptional repression would depend solely on either mononucleosomes or oligonucleosome clusters. DNA sequence accessibility would have to be increased by transient exposure of short sequences via nucleosome sliding or partial unwrapping. More sequence could be exposed by nucleosomal eviction, such as those found at steady-state in nucleosome-depleted regions. Our data, in combination with 3C suggests that yeast chromatin is best characterized as polymer-melt like, with small oligonucleosome clusters that do not pack into regular structures (Fig. 6C). The local compaction of coding regions could be a mechanism that suppresses aberrant transcriptional initiation (Struhl, 2007).

## Materials and methods

Yeast strains used in this study:

| Strain | Genotype | Source |
| --- | --- | --- |
| US1363 | *MATa bar1(unmarked) his3 leu2 trp1 ura3* | Surana lab |
| US1375 | *MATa ura3 his3 cdc20D: LEU2 GAL-CDC20::TRP1* | Surana lab |

### Chicken erythrocyte chromatin preparation

Fresh chicken blood was purchased from Nippon Bio-Test Laboratories Inc. (Japan). Erythrocyte nuclei were prepared as before (Maeshima et al., 2014a). Chromatin isolation was carried out as described by Ura and Kaneda with some modifications (Ura and Kaneda, 2001). The nuclei (equivalent to ~2 mg of DNA) in Nuclei isolation buffer (10 mM Tris-HCl pH 7.5, 1.5 mM MgCl_2_, 1.0 mM CaCl_2_, 0.25 M sucrose, 0.1 mM PMSF) were digested with 50 units of micrococcal nuclease (Worthington) at 35°C for 2 min. The reaction was stopped by addition of EGTA to 2 mM final concentration. After washing with the Nuclei isolation buffer, the nuclei were lysed with Lysis buffer (10 mM Tris-HCl pH 7.5, 5 mM EDTA, 0.1 mM PMSF). The lysate was dialyzed against Dialysis buffer (10 mM Tris-HCl pH 7.5, 0.1 mM EDTA, 0.1 mM PMSF) at 4°C overnight. The dialyzed lysate was centrifuged at 1000xG at 4°C for 5 minutes. The supernatant was recovered and used as a purified chromatin fraction. The purity and integrity of the chromatin protein components were verified by SDS-PAGE (Fig. S1A). To examine average DNA length of the purified chromatin, DNA was isolated from the chromatin fraction and electrophoresed in a 0.7% agarose gel (Fig. S1B).

### *S. cerevisiae* cell culture

Cell culture (Lim et al., 1996), G1 arrest (Yeong et al., 2000), and metaphase arrest (Liang et al., 2012) were performed as previously reported. Wild-type strain US1363 was grown in yeast-extract peptone medium (YEP) with 2 % glucose in a 24 °C water bath overnight. When the optical density at 600 nm reached ~0.5, α-factor was added to the medium to a final concentration of 0.8 μg/ml. After 3 h, the majority of cells were synchronized at G1. *cdc20Δ GAL-CDC20* strain US1375 was incubated in YEP supplemented with 2 % raffinose and 2 % galactose in a 24 °C water bath overnight and synchronized at G1 in the same way. Then the culture was filtered and washed extensively to remove α-factor and subsequently incubated in fresh YEP-glucose medium to inhibit CDC20 transcription. After 3.5 h, the majority of cells were arrested with long spindles, confirming that the *cdc20* was indeed depleted and cells were synchronized at metaphase (Yeong et al., 2000).

### Fluorescence microscopy

Microtubules were stained as before (Lim et al., 1996). *S. cerevisiae* was collected by centrifuge at 13000 RPM (15871×G) for 1 min and fixed in 1 ml 0.1 M K_2_HPO_4_ pH 6.4, 3.7% formaldehyde at 4 °C overnight. Cells were then washed and resuspended in 0.2 ml 1.2 M sorb/phos/cit (1.2 M sorbitol, 0.1 M phosphate-citrate, pH 5.9). Next, the cells were spheroplasted with 20 μl glusulase (glucuronidase > 90,000 Units/mL and sulfatase > 10,000 Units/mL) and 2 μl 10 mg/ml lyticase at 30 °C for 75 min. Then the cells were washed and resuspended in 1.2 M sorb/phos/cit. 4 μl of the sample was added to 30-well slide pre-treated with 0.1% poly-L-lysine (Sigma). Tubulin was stained with the rat monoclonal anti-α-tubulin YOL1/34 primary antibody (AbD Serotec MCA78G) and Alexa Fluor 594 conjugated goat anti-rat IgG secondary antibody (Invitrogen Molecular Probes A11007). DNA was counterstained with Vectashield-DAPI (Vector Laboratories, Inc.). The cells were imaged using a Zeiss AxioImager upright motorized microscope with Plan Apochromat 100X objective equipped with EXFO 120W metal-halide illuminator. Images were recorded on a Photometrics CoolSNAP HQ2 CCD camera controlled by Metamorph v7.7.10.0 software (Universal Imaging Corporation).

### High-pressure freezing

Yeast pellet or purified chicken-erythrocyte chromatin sample was mixed with 40-kDa dextran (Sigma) to a final concentration of 20% as an extracellular cryoprotectant. The sample/dextran mixture was loaded into a copper tube (0.45 mm outer diameter and 0.3 mm inner diameter) by using a syringe-type filler device (Part 733-1, Engineering Office M. Wohlwend GmbH). The tube was sealed at one end and high-pressure frozen using an HPF Compact 01 machine (Engineering Office M. Wohlwend GmbH). Once frozen, the tube was stored in liquid nitrogen.

### Preparation of fixed yeast for vitreous sections

Wild-type yeast (US1363, 50ml) were grown to OD_600_=0.36 in a shaker at 200 RPM at 24°C. The cells were then fixed by addition of 4.41 ml 37% formaldehyde (final concentration 3%) and incubated with shaking at 200 RPM for 15min at 30°C. Cells were pelleted by centrifugation at 4600 RPM (1987×G) for 2 min at 4°C. The supernatant was removed and the cells were resuspended in 1 ml of YEPD. The cells were washed a second time by pelleting in at 4600 RPM (1987×G) for 2min. Supernatant was removed and dextran (in YEPD medium) was added to a final concentration of 20% as extracellular cryoprotectant. The cells were then quick-spun to 3,000 RPM (845xG) to remove bubbles and then immediately high-pressure frozen as above.

### Vitreous sectioning

Vitreous sections were cut using the strategy proposed by Ladinsky (Ladinsky, 2010). Frozen-hydrated samples were cut using a Leica UC7/FC7 cryo-ultramicrotome (Leica Microsystems, Vienna, Austria) at −150 °C. First a 100 × 100 × 30 μm mesa-shaped block was trimmed using a Trimtool 20 diamond blade (Diatome). Sections were then cut using a Cryo 25° diamond knife (Diatome). The nominal thickness was set to 50–100 nm and cutting speed 1 mm/s. A customized micromanipulator (Narishige, MN-151S Model EDMS12-260) was used to control the cryo-ribbon. A “Crion” electrostatic charging device was operated in “discharge” mode to prevent the sections from sticking to the diamond blade (Pierson et al., 2010). Once the ribbon was 2–3 mm long, it was transferred onto an EM grid (Protochips C-flat or EMS continuous carbon grid Cat# CF-200-CU-50) to which 10-nm gold fiducials (BBI Cat# EM.GC10) were already added. This transfer was initiated by operating the Crion in “charge” mode. Subsequently, the ribbon was physically pressed with a 10-mm laser window glass (Edmund Optics stock # 65-855) to ensure firm attachment. The grid was stored in liquid nitrogen until imaging.

### Plunge freezing

Plunge freezing was done using Vitrobot MARK IV (FEI, Eindhoven) operated at 20 °C with 100% humidity. Purified chromatin sample was mixed with 10-nm gold fiducials (BBI, as above) and 3 μl of this mixture was applied onto freshly glow-discharged EM grids with hole diameters and spacings appropriate for the sample type (see Table S2). The grid was blotted once and then plunged into a liquid propane-ethane mixture (Tivol et al., 2008).

### Electron cryotomography

Tilt series were collected using Leginon (Suloway et al., 2009) or FEI TOMO 3 & 4 on a Titan Krios cryo-TEM (FEI, Eindhoven) operating at 300 KeV. Tomography data was recorded on either Falcon I or Falcon II direct-detection cameras. Imaging parameters for each sample type are listed in Table S2. Image alignment, CTF correction, 3-D reconstruction and visualization were done using IMOD software package (Kremer et al., 1996; Mastronarde, 1997; Xiong et al., 2009). Default settings were used except that the low pass filter cutoff was set to 0.3.

### Fourier analysis

Tomographic slices were imported into ImageJ 1.49v (Schneider et al., 2012). The Fourier transform was calculated using the FFT function. The radial plot of the FT was generated using Radial Profile Angle plugin (http://rsbweb.nih.gov/ij/plugins/radial-profile-ext.html). The plot values (radius and normalized intensities) were saved. Radius values in pixels were converted to values in real space based on the pixel size at the specimen level. The plot was generated using Excel (Microsoft Office, version 14.1.0) and saved as an image.

### Three-dimensional density visualization

Isosurface rendering was done with the UCSF Chimera package (Pettersen et al., 2004). Subtomograms were normalized to a mean of 0 and standard deviation of 1 using EMAN2 (Tang et al., 2007), and the contour level was set to 1.5σ. Isosurface densities smaller than 6 nm were removed using the “hide dust” function.

### Template matching

Template matching was done using PEET, which accounts for the tomographic missing wedge (Heumann, 2016; Nicastro et al., 2006). Oligonucleosome reference models were generated using Bsoft (Heymann and Belnap, 2007). To minimize the effects of adjacent densities in the highly crowded intranuclear environment, we applied either a squat or elongated cylindrical mask (depending on the aspect ratio of the reference) around the oligonucleosome reference. Overlapping hits were automatically subjected to duplicate removal at the end of template matching. The top 10% of hits were visually inspected to remove the remaining spurious hits.

## Figures & media

All figures were composited in Adobe Photoshop or Adobe Illustrator; movie S1 was assembled and rendered with Adobe Premiere Pro (Adobe Systems, Inc., San Jose).

## Data sharing

An example tomogram (corresponding to Fig. 2B) has been deposited at the EMDB (EMD-8157). Tilt series (raw) data and IMOD reconstruction parameters of all samples presented in this paper have been made publicly accessible in the EMPIAR online database: (EMPIAR-10076) (Iudin et al., 2016). Details of the corresponding figure and sample are summarized in Table S1.

## Acknowledgements

We thank the CBIS cryo-EM staff for support and especially Mdm. Loy for microtomy training; Mark Ladinsky for cryomicrotomy advice; and Benoît Zuber, Danny Studer, and Dmitri Vanhecke for cryomicrotomy training at the Institute of Anatomy of the University of Bern. We thank John Heumann and David Mastronarde for advice on PEET and IMOD, and Matthijn Vos for Titan Krios training. We thank our CBIS colleagues and the Gan group for feedback on the manuscript. CC and LG were supported by NUS startups R-154-000-515-133, R-154-000-524-651, and D-E12-303-154-217, with equipment support from NUS YIA R-154-000-558-133 and MOE T2 R-154-000-624-112. HHL and US were funded by the Biomedical Research Council of A*STAR (Agency of Science Technology and Research), Singapore. ST and KM were supported by JST CREST grant and MEXT KAKENHI grant (23115005).

## Abbreviations list

Yeast, *S. cerevisiae*; Cryo-EM, Electron cryomicroscopy; Cryo-ET, Electron cryotomography; 3C, chromatin conformation capture.

---

**Figure S1.**
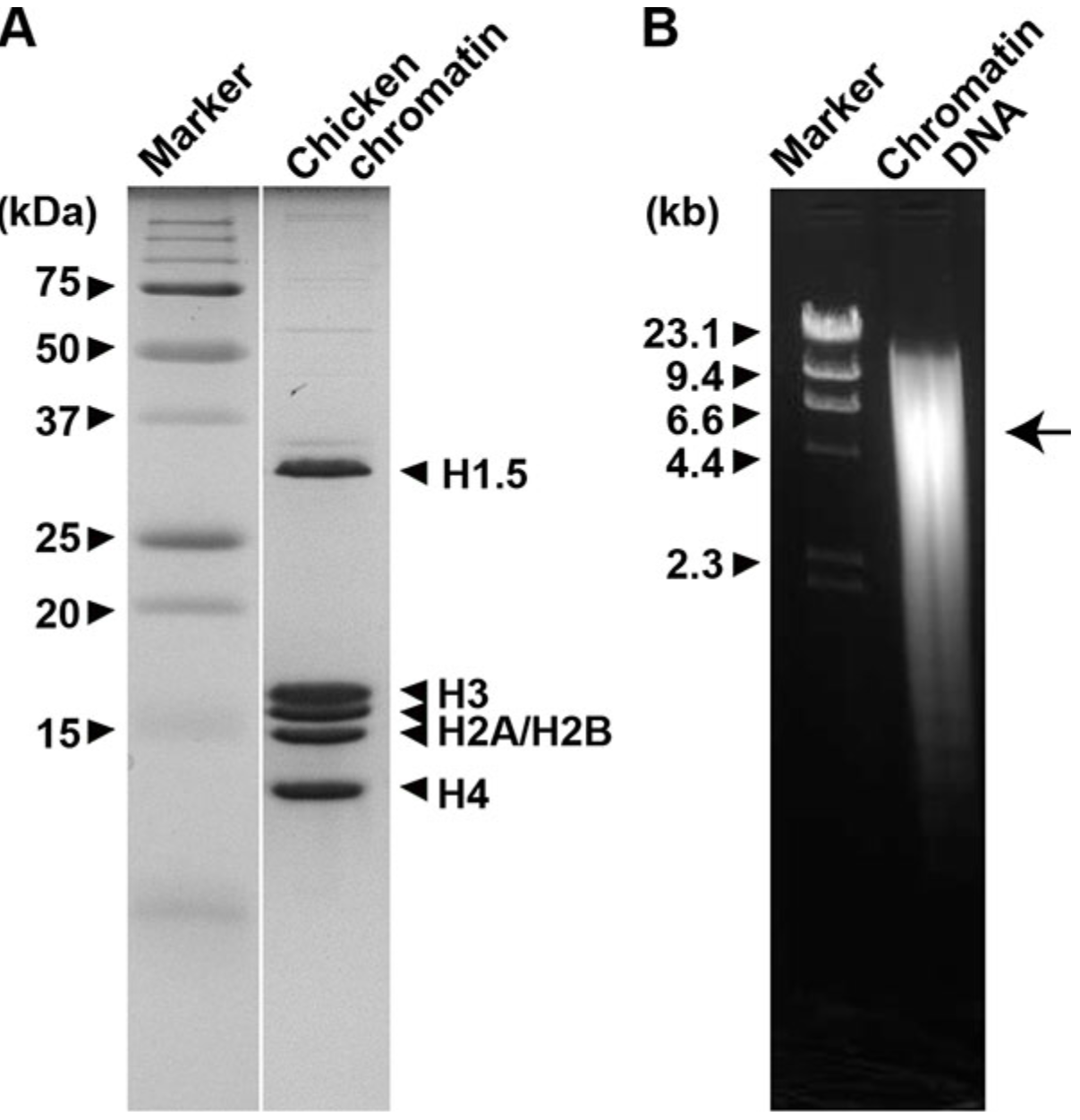
Isolation of chicken erythrocyte chromatin. **(A)** SDS-PAGE of purified chromatin. Nucleosome core histones H2A/H2B, H3 and H4 as well as linker histone H1.5 are present in the sample. **(B)** Agarose gel of isolated chromatin DNA. Average DNA length is ~5 kb (arrow).

**Figure S2.**
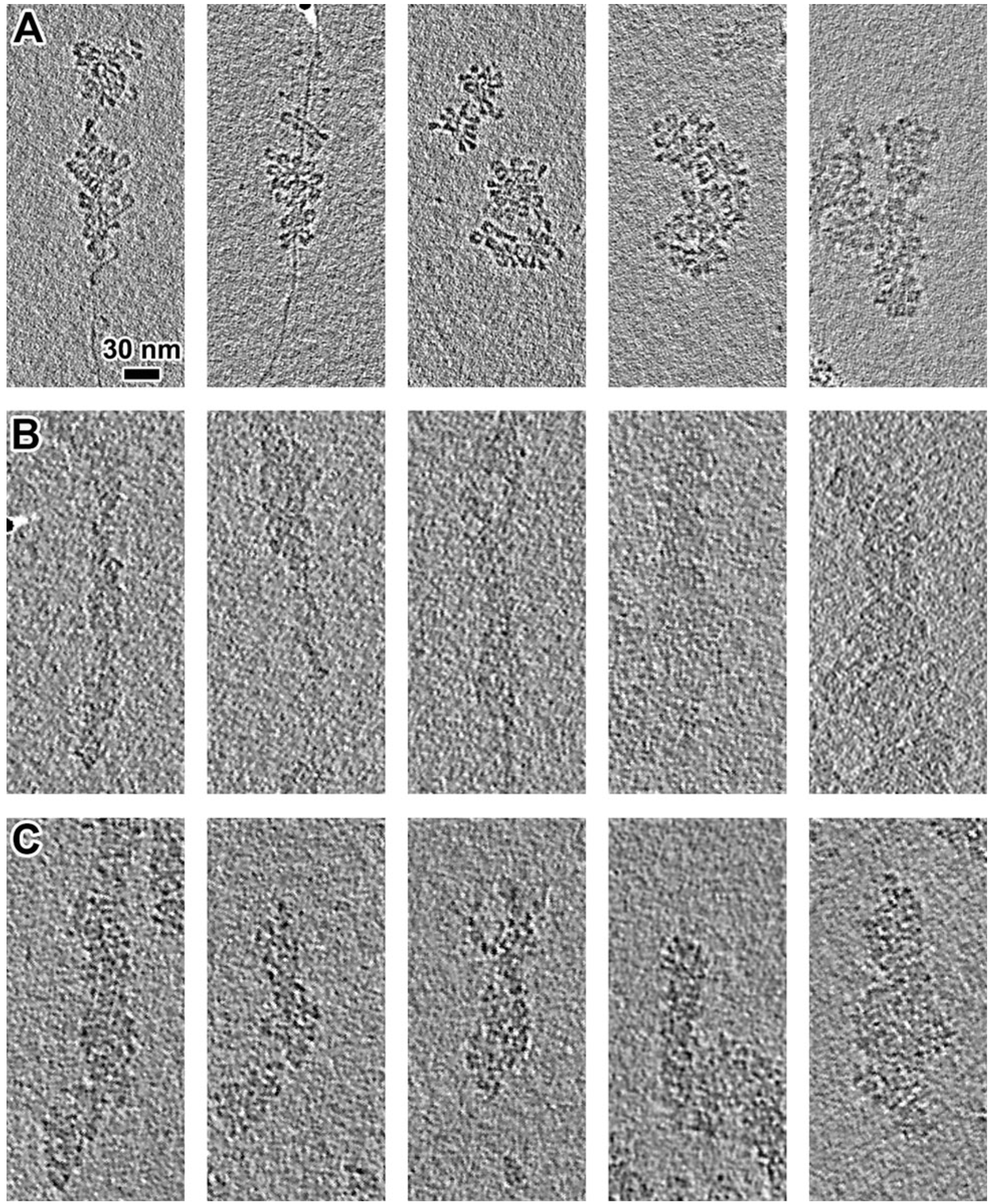
Chromatin fibers are recognizable regardless of cryo-EM sample-preparation method. Row **(A)**: plunge-frozen in isolation buffer; row **(B)**: plunge-frozen in isolation buffer plus 20% 40-kDa dextran; row **(C)**: high-pressure frozen in isolation buffer plus 20% dextran, then cryosectioned. Note that the addition of dextran substantially lowers the image contrast.

**Figure S3.**
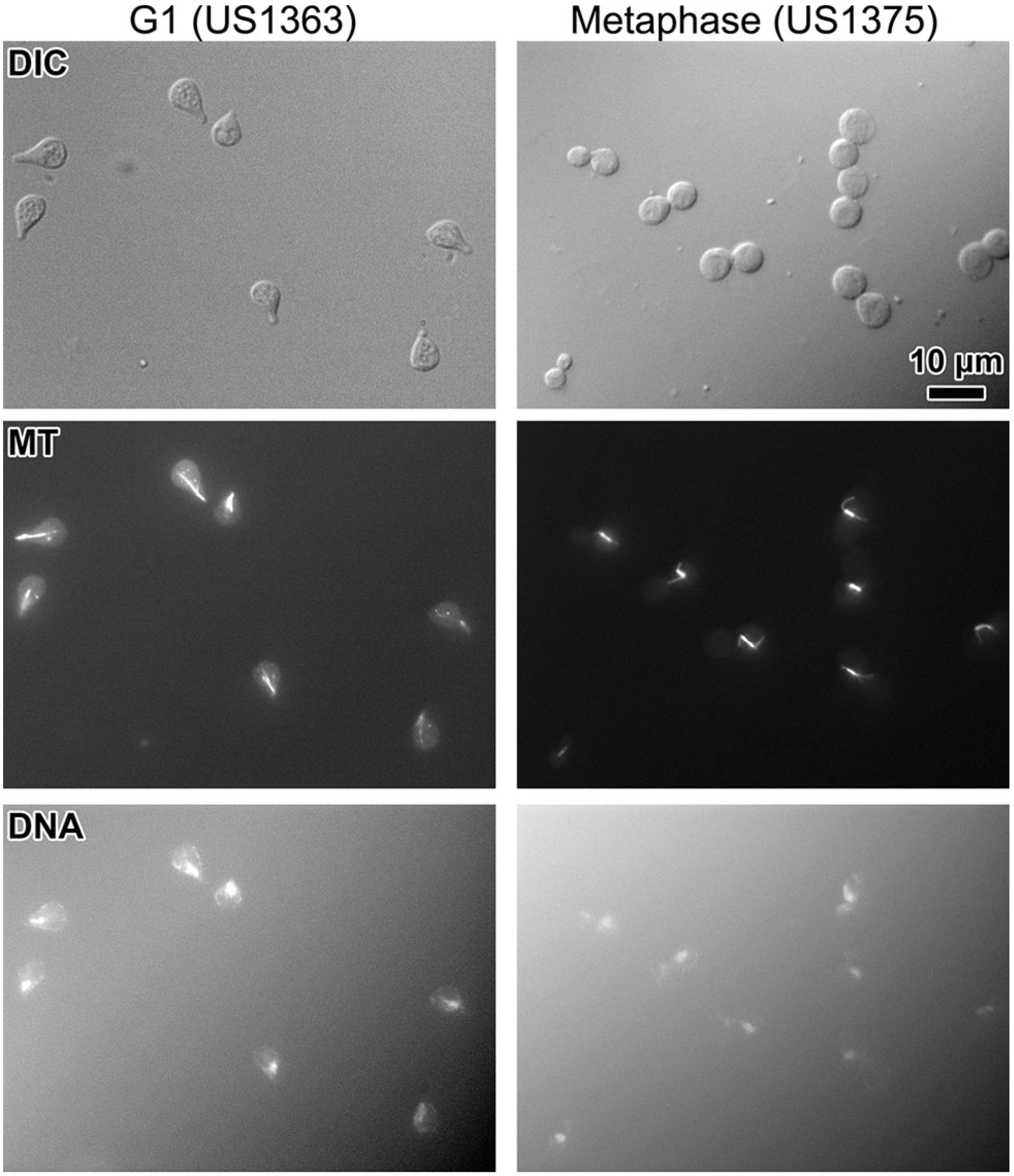
Synchronization of *S. cerevisiae*. US1363 cells are arrested at G1 and show the characteristic “Shmoo” morphology (left column) when incubated with α-factor; *cdc20Δ GAL-CDC20* (US1375) mutants are arrested at metaphase in presence of glucose. These metaphase cells have a characteristic large-bud morphology and a short, bar-shaped spindle (right column).

**Figure S4A.**
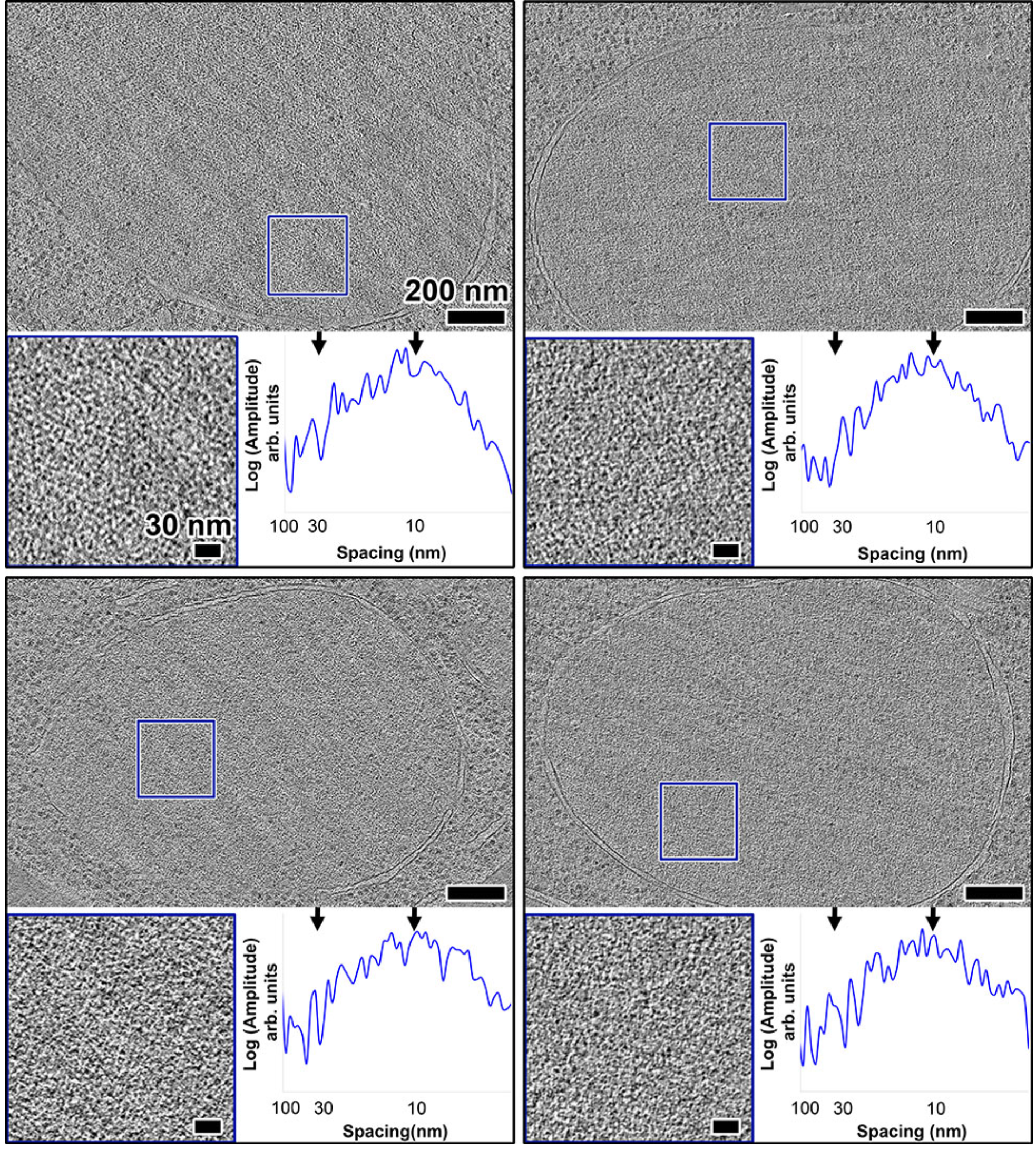
Additional examples of G1-arrested cells. Tomographic slices (30-nm thick) of four more G1 cells. The lower left subpanel is a 3-fold enlargement of the area boxed in blue. The lower right subpanel shows a rotationally averaged power spectrum of this area. Arrows indicate 30- and 10-nm spacings.

**Figure S4B.**
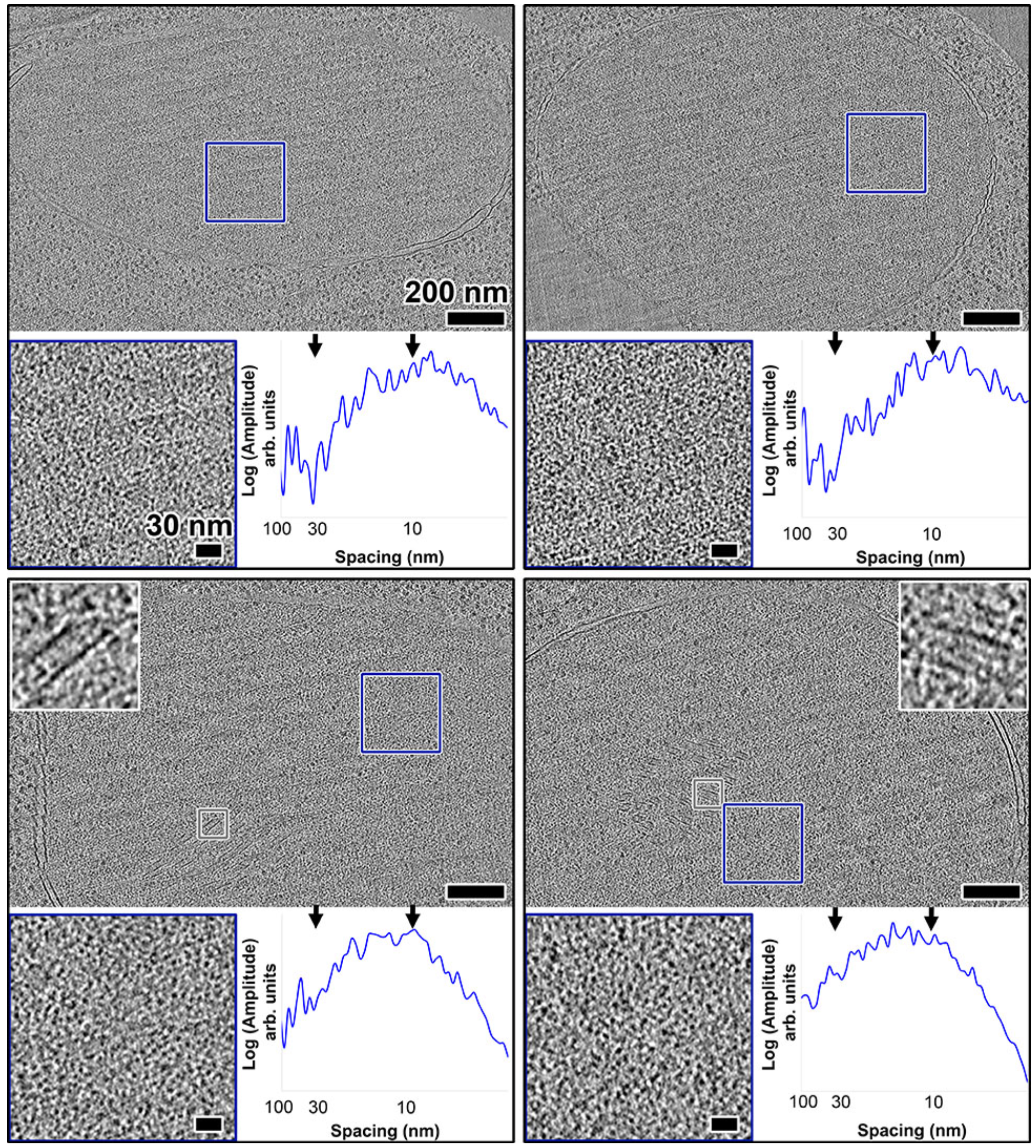
Additional examples of metaphase-arrested cells. Tomographic slices (30-nm thick) of four more metaphase cells. The lower left subpanel is a 3-fold enlargement of the area boxed in blue. The lower right subpanel shows a rotationally averaged power spectrum of this area. Arrows indicate 30- and 10-nm spacings. For the two cells in the lower panels, the upper 792 subpanels show spindle microtubules, boxed in grey and enlarged 5-fold.

**Figure S4C.**
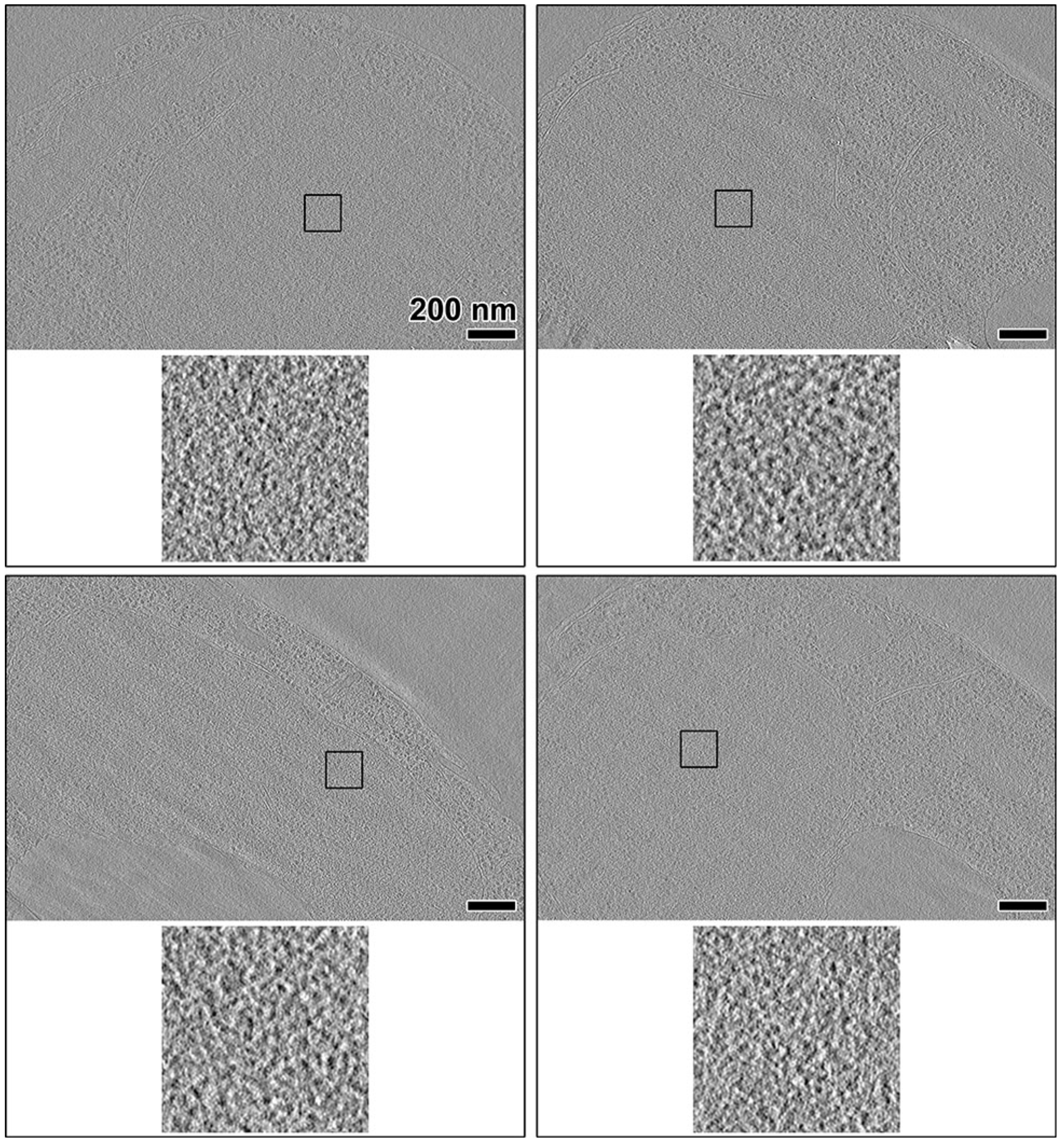
Additional examples of formaldehyde-fixed cells. Tomographic slices (10-nm thick) of four more wild-type cells, fixed in formaldehyde. A region inside each nucleus is boxed out and enlarged 6-fold in the lower subpanel.

**Figure S5.**
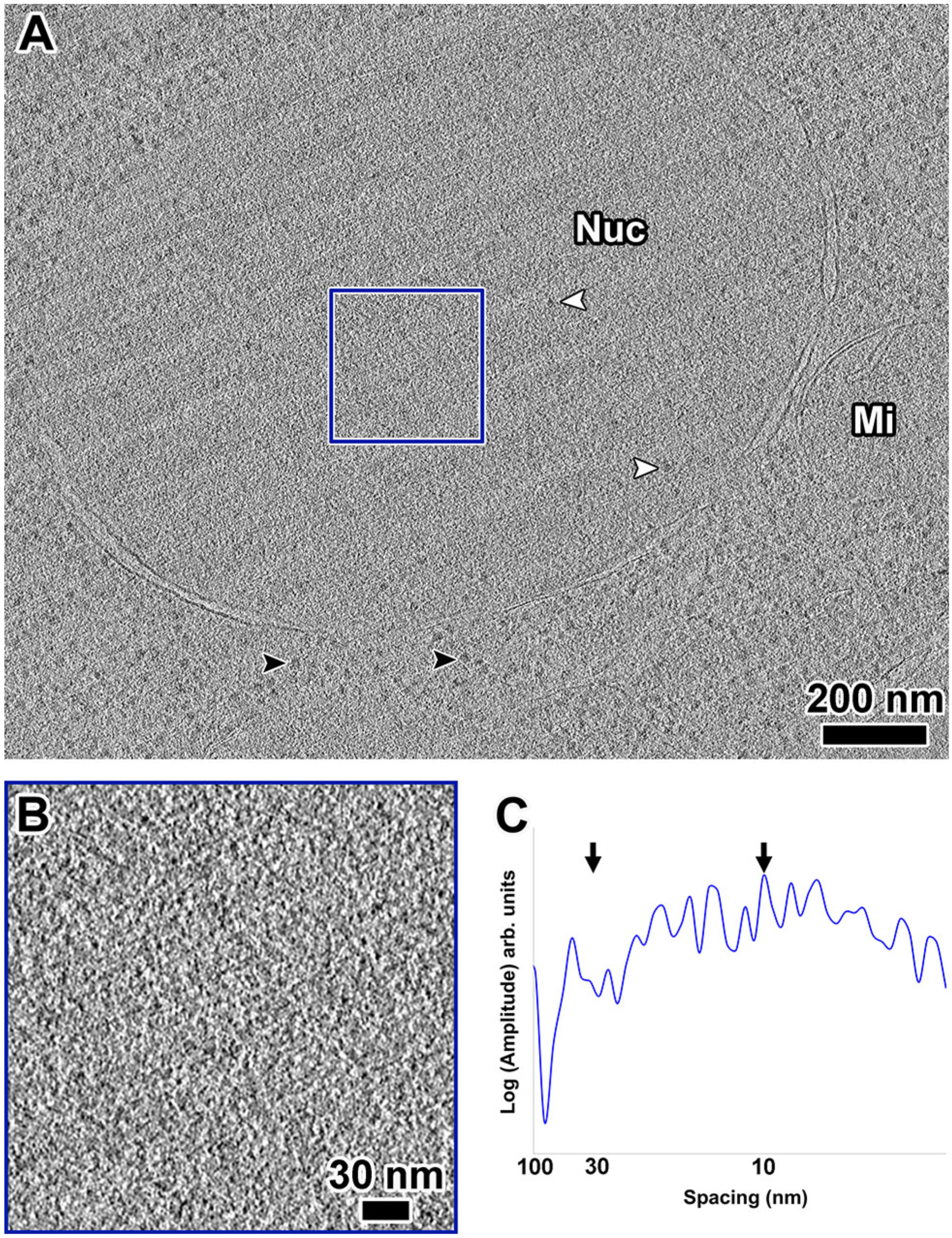
Chromatin fiber signals are absent regardless of imaging conditions. (A) Tomographic slice (30-nm thick) of a nucleus in a G1-arrested cell that was imaged much closer to focus (−3 μm) than the other cells presented in this paper. The nucleus (Nuc) and a mitochondrion (Mi) are labeled. White arrows: intranuclear ribosome-sized densities; black arrows: cytoplasmic ribosome densities. (B) A 3-fold enlargement of the intranuclear position boxed in panel (A). (C) Rotationally averaged power spectrum of (B). Arrows point to 30- and 10-nm spacings.

**Table S1.**
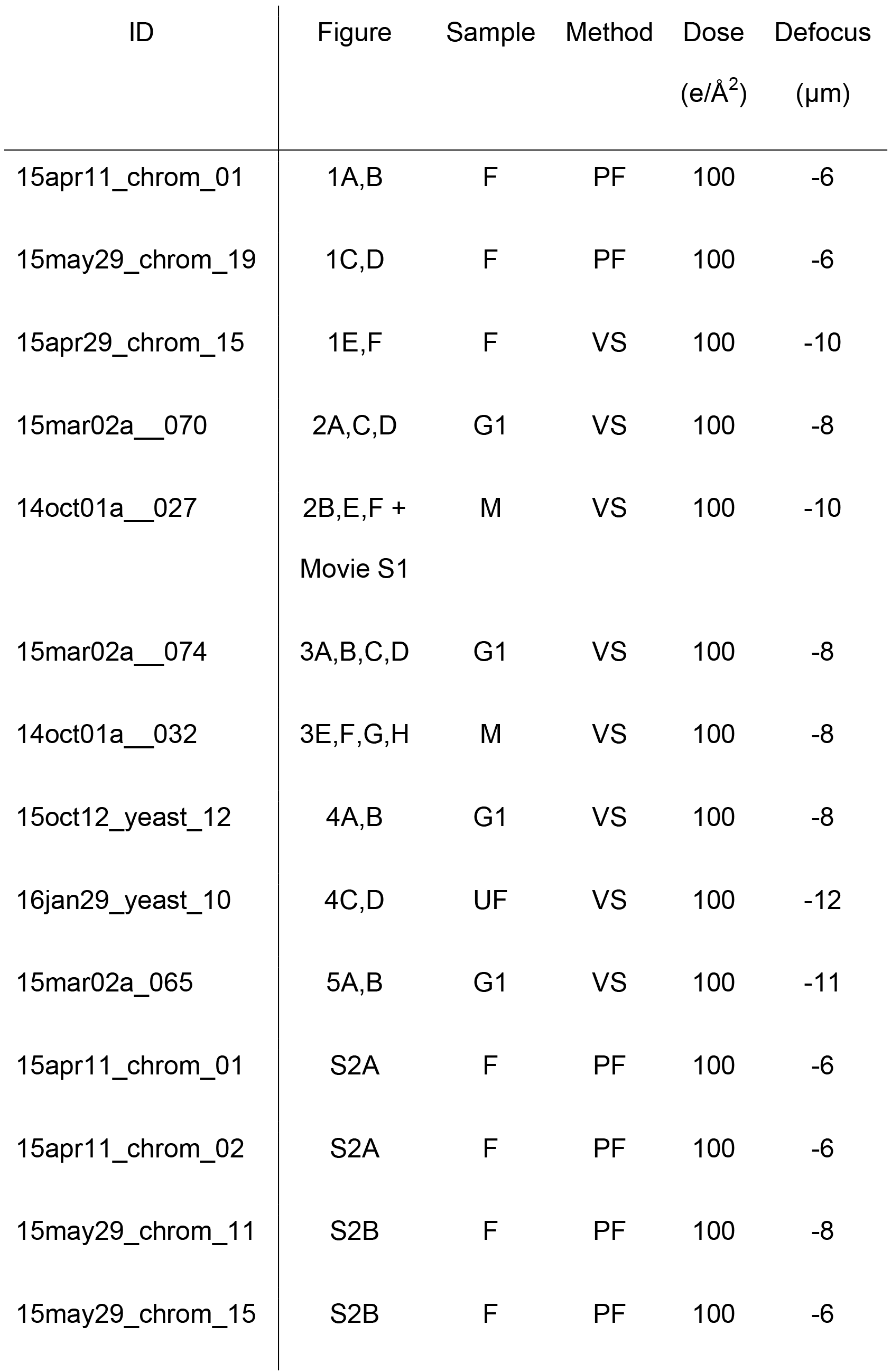
Tomogram details.

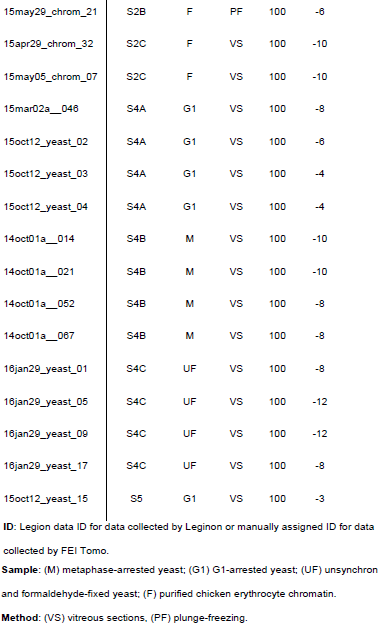

**Table S2.** Cryotomography imaging conditions.

| Sample | Purified chromatin | *S. cerevisiae* cells |
| --- | --- | --- |
| EM grid | CF-22-2C-T (Protochips) | CF-42-2C-T (Protochips) /<br>continuous carbon (EMS) |
| Microscope | Titan Krios |  |
| Energy | 300 keV |  |
| Gun type | FEG |  |
| Camera | Falcon II Direct Detector | Falcon I or II Direct Detector |
| Tomography software | FEI TOMO 3 and 4 | Leginon |
| Calibrated magnification | 23986 | 15678 / 19167 |
| Calibrated pixel size | 5.84 Å | 8.93 / 7.47 Å |
| Defocus | -6 to -8 µm | -3 to -15 µm |
| Cumulative dose | 80 - 100 electrons / Å<sup>2</sup> |  |
| Dose fractionation | 1/cos |  |
| Tilt range | ±60° |  |
| Tilt increment | 2° |  |

